# protPheMut: An Interpretable Machine Learning Tool for Classification of Cancer and Neurodevelopmental Disorders in Human Missense Variants

**DOI:** 10.1101/2025.01.06.631365

**Authors:** Jingran Wang, Miao Yang, Chang Zong, Gennady Verkhivker, Xiao Fei, Guang Hu

**Affiliations:** MOE Key Laboratory of Geriatric Diseases and Immunology, Suzhou Key Laboratory of Pathogen Bioscience and Anti-Infective Medicine, Department of Bioinformatics and Computational Biology, School of Life Sciences, Suzhou Medical College of Soochow University, Suzhou 215213, China; Department of Computational and Data Sciences, Chapman University, One University Drive, Orange, California, United States of America; Department of Biomedical and Pharmaceutical Sciences, Chapman University Pharmacy School 9401 Jeronimo Rd, Irvine, California, United States of America; Jiangsu Province Engineering Research Center of Precision Diagnostics and Therapeutics Development, Soochow University, Suzhou 215123, China; Key Laboratory of Alkene-carbon Fibres-based Technology & Application for Detection of Major Infectious Diseases, Soochow University, Suzhou 215123, China; Jiangsu Key Laboratory of Infection and Immunity, Soochow University, Suzhou 215123, China

## Abstract

**Motivation:** Recent advances in human genomics have revealed that missense mutations in a single protein can lead to distinctly different phenotypes. In particular, some mutations in oncoproteins like Ras, MEK, PI3K, PTEN, and SHP2 are linked various cancers and Neurodevelopmental Disorders (NDDs). While numerous tools exist for predicting the pathogenicity of missense mutations, linking these variants to certain phenotypes remains a major challenge, particularly in the context of personalized medicine.

**Results:** To fill this gap, we developed protPheMut (Protein Phenotypic Mutations Analyzer), leveraging multiple interpretable machine learning methods and integrate diverse biophysics and network dynamics-based signatures, for the prediction of mutations of the same protein can promote cancer, or NDDs. We illustrate the utility of protPheMut in phenotypes (cancer/NDDs) prediction by the mutation analysis of two protein cases, that are PI3Kα and PTEN. Compared to seven other predictive tools, protPheMut demonstrated exceptional accuracy in forecasting phenotypic effects, achieving an AUROC of 0.8501 for PI3Kα mutations related to cancer and Cowden syndrome. For multi-phenotypes prediction of PTEN mutations related to cancer, PHTS, and HCPS, protPheMut achieved an AUC of 0.9349 through micro-averaging. Using SHAP model explanations, we gained insights into the mechanisms driving phenotype formation. A userfriendly website deployment is also provided.

**Availability:** Source code and data are available at https://github.com/Spencer-JRWang/protPheMut. We also provide a user-friendly website at http://netprotlab.com/protPheMut.

**Supplementary information:** Supplementary data are available at *Bioinformatics* online.

**Graphical Abstract:** 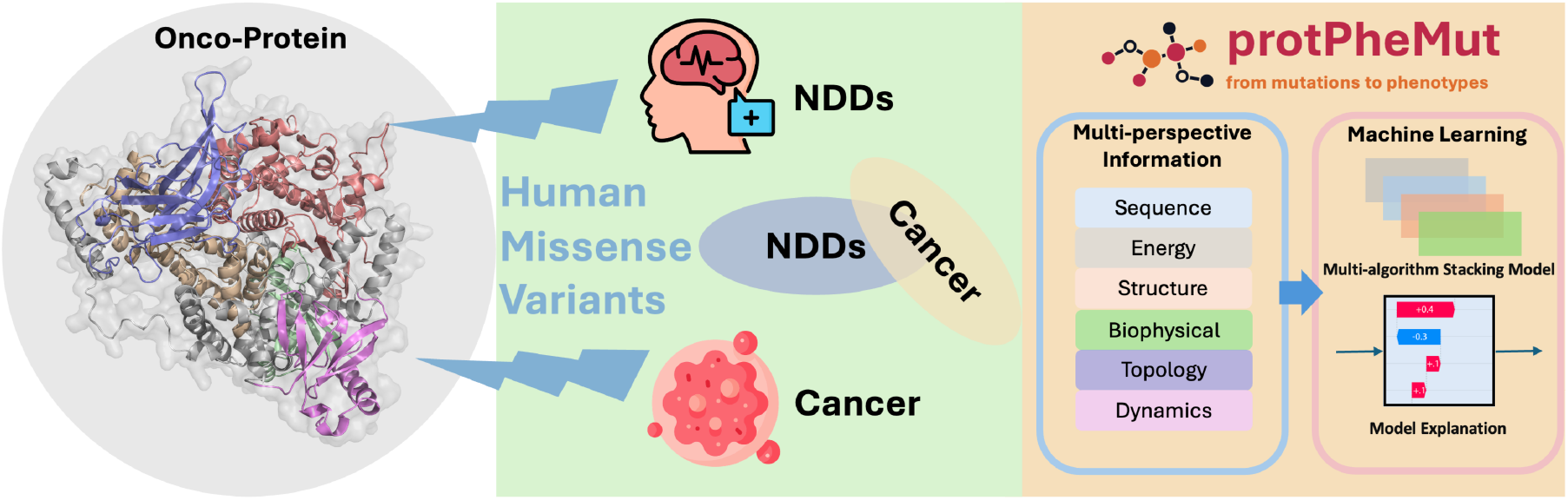

## 1 Introduction

Neurodevelopmental disorders (NDDs) encompass a range of developmental abnormalities, such as RASopathies, Cowden syndrome, autism spectrum disorder (ASD), and PTEN Hamartoma Tumor Syndrome (PHTS). Numerous studies have demonstrated a strong correlation between NDD and cancer development, showing that individuals with NDD are at a higher risk of developing cancer (Li, et al., 2020; Nussinov, et al., 2022; Nussinov, et al., 2024). NDDs and cancer share key proteins, pathways, and mutations, and variations in pathway signaling strengths have been identified as critical factors of different phenotypes (Nussinov, et al., 2023; Yavuz, et al., 2023). Mutations in oncogenic proteins such as Ras, MEK, PI3K, phosphatase and tensin homolog (PTEN), and SHP2 are linked to both cancer and NDDs (Liu, et al., 2023). Recent research has explored the connections between these conditions, covering topics such as RASopathies (e.g., neurofibromatosis type 1, Noonan syndrome, Costello syndrome) (Gross, et al., 2020), PIK3CA-related overgrowth spectrum (Venot, et al., 2018), Cowden syndrome (Orloff, et al., 2013), ASD (Yehia, et al., 2020), as well as genetically engineered mouse models. There is growing evidence linking developmental signaling pathways to aggressive central nervous system tumors (Nussinov, et al., 2022). Despite this progress, a key question remains: why do mutations in the same protein can lead to vastly different phenotypic and clinical outcomes, particularly in cancer and NDDs.

Predicting the effects of missense mutations has become a key focus in computational medicine (Cheng, et al., 2020). Traditional methods, such as molecular dynamics (MD) simulations, have shown success in assessing the impact of individual mutations but require significant computational resources and time (Elia Venanzi, et al., 2024), limiting their feasibility for large-scale studies. The information derived from protein missense mutations is inherently multi-perspective, and accurate prediction of their effects requires the integration of various dimensions of data. Recent tools based on sequence conservation and co-evolution of amino acid residues, such as PolyPhen2 (Adzhubei, et al., 2013) and EVMutation (Hopf, et al., 2017), have been widely applied to predict the pathogenicity of human variants. Similarly, MutPred2 successfully integrates multi-perspective data using ensemble neural networks to predict detailed mutation effects (Pejaver, et al., 2020). Moreover, a recent study added local environment and interaction features and successfully classify PTEN related multi-phenotypes (Portelli, et al., 2021). With the advent of AI, tools like AlphaMissense, based on AlphaFold2 (Jumper, et al., 2021), are achieving high-precision predictions for a broad range of human protein missense mutations (Cheng, et al., 2023). However, current methods primarily address the pathogenicity or functional alterations caused by mutations, leaving a pressing challenge in predicting distinct phenotypic outcomes, such as cancer versus NDDs.

Proteins exist as dynamic conformational ensembles, which can act as phenotypic determinants of mutations (Nussinov, et al., 2023). Incorporating dynamics features has proven beneficial for evaluating the pathogenicity of single amino acid variants (Barozi, et al., 2024; Pacini and Lesieur, 2022). For instance, Rhapsody (Ponzoni and Bahar, 2018; Ponzoni, et al., 2020) leverages the Elastic Network Model (ENM) to analyze intra-molecular dynamics based on the topology of residue contact networks, coupled with machine learning for mutation effect predictions. In our previous works (Liang, et al., 2020; Verkhivker, et al., 2020; Zhu, et al., 2023; Zhu, et al., 2022), we demonstrated the unique utility of intrinsic dynamical features in predictive ML and AI models. In particular, by building a multi-faceted platform including features from sequence-structure-dynamics, we have characterized the allosteric effects of ALPL mutations and role of allostery in the pathogenesis of HPPs (Xiao, et al., 2022).

In this work, we introduce protPheMut, an interpretable ML-based tool that combines Gradient Boost Machine (GBM) algorithms with a stacking model to integrate diverse biophysics and network dynamics-based features. This framework enables accurate classification of missense mutations associated with cancer and NDDs while maintaining model interpretability to uncover phenotype-specific mutation characteristics. protPheMut is available as a user-friendly web tool, supporting automated retrieval of AlphaFold2-predicted structures, experimental data, or homology-modeled protein structures, along with phenotype-labeled missense mutations. By automating feature extraction, model training, and evaluation, protPheMut streamlines mutation analysis, as demonstrated in its application to PI3Kα and PTEN proteins.

## 2 Materials and methods

### 2.1 Data collection of protein structure and mutations

Protein structures were derived from AlphaFold2 prediction, accessed via the AlphaFold2 database (Varadi, et al., 2024). Pathogenic missense mutations proteins are collected from COSMIC Database (Sondka, et al., 2024), Clinvar Database (Landrum, et al., 2014), The Genome Aggregation Database (Chen, et al., 2024), UniProt database (UniProt, 2023), The Human Gene Mutation Database (Stenson, et al., 2020), and DisGeNET platform (Pinero, et al., 2020).

For PI3Kα, a total of 782 mutations were gathered: 154 mutations related to NDD-associated Cowden syndrome and 628 mutations related to cancer. Similarly, for PTEN, we collected totally 653 mutations related to three phenotypes: 70 mutations related to NDD-associated Hereditary Cancer-Predisposing Syndrome (HCPS), 150 mutations related to PTEN Hamartoma Tumor Syndrome (PHTS), and 453 mutations related to cancer. All missense mutations with their phenotype labels are listed in Supplementary Table 1.

To ensure consistency and relevance in our analysis, the collected mutations adhered to the following criteria. Firstly, the mutations must be pathogenic missense mutations and single nucleotide variants (SNVs), emphasizing those with clear molecular mechanisms and clinical significance. Secondly, mutations associated with multiple phenotypes were excluded to avoid potential confounding factors, ensuring a more straightforward interpretation of phenotype formation.

### 2.2 protPheMut framework

Figure 1 illustrates the overall flowchart of protPheMut framework, which contains mainly two parts: the feature generation module and machine learning module. The feature generation module automatically calculates a diverse set of features that capture multi-perspective information, including sequence conservation, energy and structural properties, physicochemical characteristics, network topology, and intermolecular dynamics. To be more concrete, the features generated include entropy, coevolution, conservation score, hydrophobicity, RASA (Relative Accessible Surface Area), ΔΔG (protein folding energy change), ΔBC (node betweenness centrality change), ΔC (node closeness centrality change), ΔEC (node eigenvector centrality change), ΔCC (node clustering coefficient centrality change), effectiveness, sensitivity, stiffness, MSF (Mean Square Fraction), and DFI (Dynamic Flexibility Index). The ML model performs automatic feature selection and employs multi-stacking models based on GBM and Logistic Regression. Cross-validation prediction is employed to predict a phenotypic probability (PhenoScore) for each mutation. Additionally, SHAP-based model enhances both global and local explanation. For multiple phenotypes classification, we apply One-vs-One strategy towards stacking model. A more detailed framework description can be found in Supplementary Text 1.

**Fig. 1.**
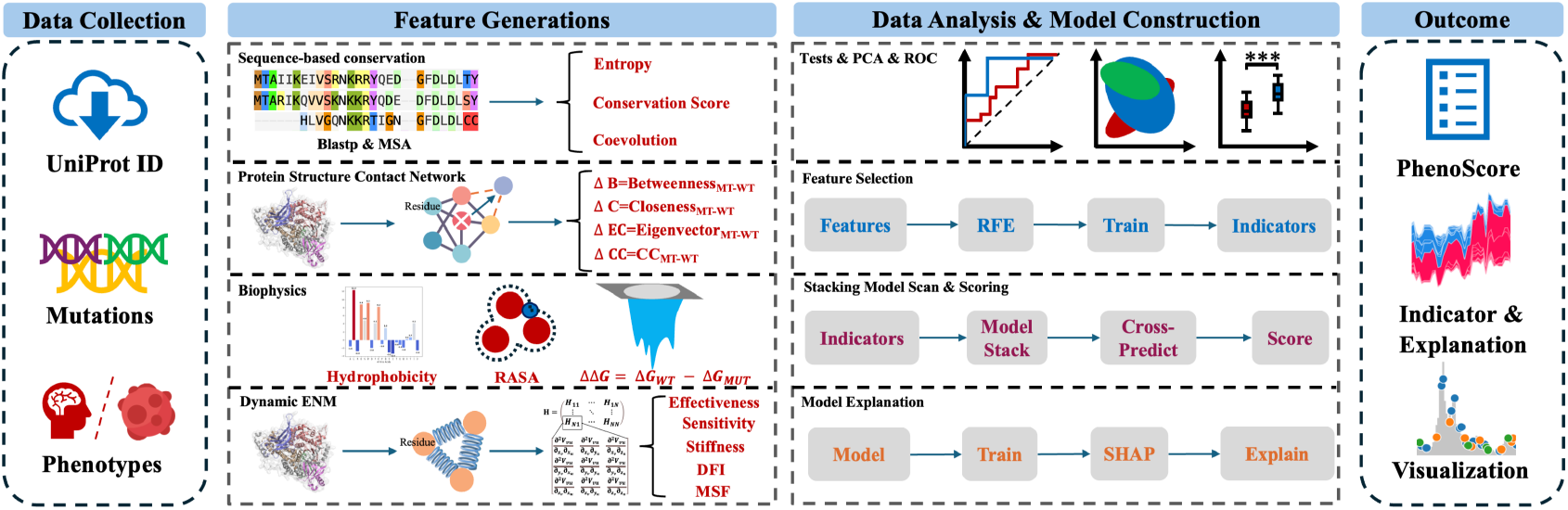
The computational workflow overview of protPheMut. protPheMut uses AlphaFold prediction structure by default, users can also upload their own pdb file, and then sequence-based features, contact energy network related features, Structure and energy related features and elastic network model-related features are integrated by machine learning model to generate PhenoScore, indicators, model interpretation and automatic visualization.

### 2.3 Model evaluation and comparison

Since no existing tool specifically addresses the phenotypic effects of protein mutations, we evaluated the performance of protPheMut by comparing it with eight widely used tools for pathogenicity prediction, including AlphaMissense (Cheng, et al., 2023), EVMutation (Hopf, et al., 2017), PolyPhen-2 (Adzhubei, et al., 2013), Rhapsody (Ponzoni, et al., 2020), MutPred2 (Pejaver, et al., 2020), FATHMM (Shihab, et al., 2013), and SIFT (Sim, et al., 2012). We used various metrics to assess the predictive performance of the models, including accuracy, precision, recall, specificity, sensitivity, F1 score, AUROC (Area Under the Receiver Operating Characteristic Curve) and AUPRC (Area Under the Precision-Recall Curve).

### 2.4 Webpage deployment

To optimize the application of our methodological framework, we developed a web-based platform based on HTML5, JavaScript, jQuery, and PHP. This platform allows users to input protein structures along with mutations that are annotated with phenotype labels. Upon submission, the protPheMut framework is automatically executed to analyze the input data and generate results, streamlining the study of genotype-phenotype relationships.

## 3 Results

Oncoproteins related to NDDs are involved in several major pathways. To demonstrate the utility of our method for the classification of cancer and NDDs, we focus on two oncoproteins, PI3Kα and PTEN, involved in the PI3K/AKT/mTOR pathway.

### 3.1 Application to PI3Kα mutations in classifying Cowden syndrome and cancer

Figure 2a illustrates the distribution of PI3Kα mutations along the protein sequence. PI3Kα consists of five main domains: PI3K_p85B, PI3K_rbd, PI3K_C2, PI3Ka, and PI3_PI4_kinase. Of the collected mutations, 154 (19.7%) were linked to Cowden syndrome, and 629 (80.3%) were associated with cancer, as shown in Figure 2b. These mutations were distributed across the structural domains of PI3Kα. while Figures 2c, d provide structural mapping of PI3Kα mutations associated with cancer and Cowden syndrome (Orloff, et al., 2013).

**Fig. 2.**
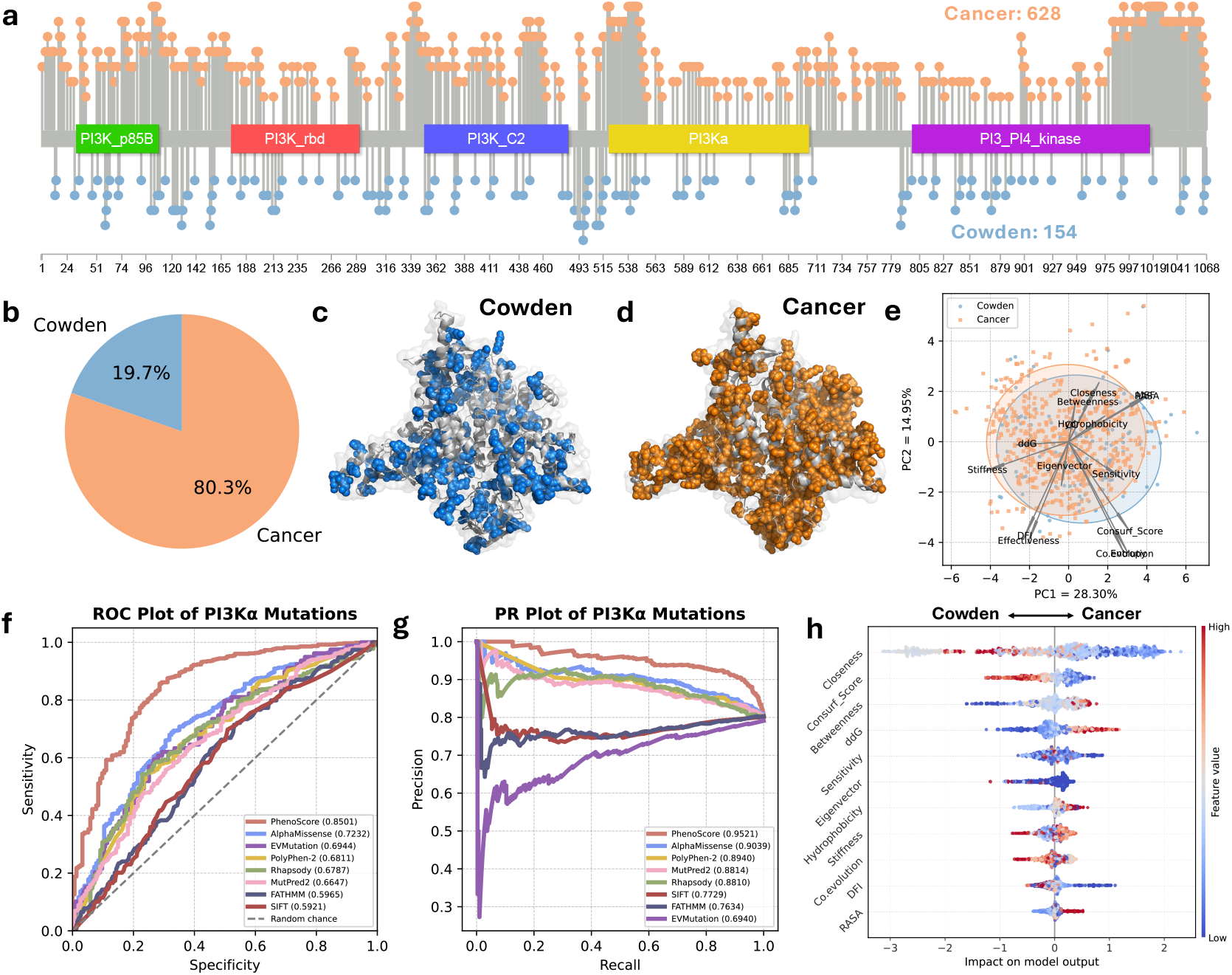
protPheMut classify mutations’ binary phenotypic effects. a) The lollipop plot shows the distribution of missense mutations on PI3Kα protein and mutation’s phenotypes. b) The pie chart shows the Cowden syndrome and cancer variants in dataset. c) The distribution of mutations related to Cowden syndrome in PI3Kα structure. d) The distribution of mutations related to cancer in PI3Kα structure. e) The PCA plot of Cowden and cancer mutations with features’ direction. f) The comparison of PhenoScore generated from 10-fold straight cross-validation with 8 other predicted tools in the whole dataset. g) Precision-recall curve of PhenoScore and 8 other predicted tools in the whole dataset. h) Beeswarm plot shows the global and local interpretation of the model using SHAP. Features higher up on the plot indicate greater importance. Red and blue represent the magnitude of the feature values, and their distribution along the X-axis reflects the positive or negative contribution of each feature on predicting single mutation.

The protPheMut tool was employed to automatically calculate total 15 features of these mutations. Principal Component Analysis (PCA) of these features (Figure 2e) revealed distinct yet overlapping distributions of cancer and Cowden syndrome samples. The first two principal components (PC1 and PC2) accounted for 28.30% and 14.95% of the variance, respectively, with arrows indicating the contribution of each feature.

Using LightGBM-based recursive feature elimination (RFE), we identified 11 key features that effectively differentiated PI3Kα mutations linked to Cowden syndrome and cancer. These features included sequence-level metrics (conservation score and coevolution), energy change (ΔΔG), structure-level properties (RASA), network centrality metrics (ΔBC, ΔC, and ΔEC), biophysical changes (hydrophobicity), and dynamic properties (sensitivity, stiffness, and DFI). We tested all combinations of 4 gradient boosting models (Gradient Boost, XGBoost, LightGBM, and CatBoost), and integrated the predictive scores using logistic regression. The stacking model built with 2 of the models (Gradient Boost and CatBoost) achieved the best performance in distinguishing Cowden syndrome from cancer, with a PhenoScore AUC of 0.8501 and an AUPRC of 0.9521 using 10-fold straight cross-validation (Figure 2f, g). Current tools primarily predict mutation pathogenicity but are not optimized for phenotypic classification. Compared to protPheMut, these tools, including AlphaMissense, EVMutation, PolyPhen-2, Rhapsody, MutPred2, SIFT, and FATHMM, had limited success in distinguishing between Cowden syndrome and cancer. Among them, AlphaMissense showed the best performance, with an AUC of 0.7232 and an AUPRC of 0.9039. These results highlight that pathogenicity scores alone are insufficient for phenotypic classification, whereas protPheMut outperformed all benchmarks.

Figure 2h illustrates the model interpretation, with the most important features listed at the top. Based on model interpretation, it highlights the interplay of bio-graph, dynamic, energy and evolutionary information in distinguishing between cancer and Cowden syndrome, with the magnitude and direction of each feature’s contribution shaping the final prediction. To be more concrete, compared to Cowden syndrome mutations, cancer mutations show decrease in ΔCC, conservation score, sensitivity and show increase in ΔBC and ΔΔG. More detailed analysis of PI3Kα can be found at Supplementary Text S2.

### 3.2 Application to PTEN mutations in classifying PHTS, HCPS, and cancer

Figure 3a shows the distribution of PTEN mutations along the protein sequence. PTEN consists of two main domains: the Tc-R-P domain and the C2 domain. Figure 3b shows the properties of PTEN mutations, we collected a total of 70 mutations related to HCPS (10.4%), 150 mutations related to PHTS (80.4%), and 454 mutations related to cancer (67.4%), distributed across the structural domains of PTEN. Figures 3c-e provide structural mappings of PTEN mutations associated with PTHS, HCPS, and cancer (Portelli, et al., 2021; Yehia, et al., 2020).

**Fig. 3.**
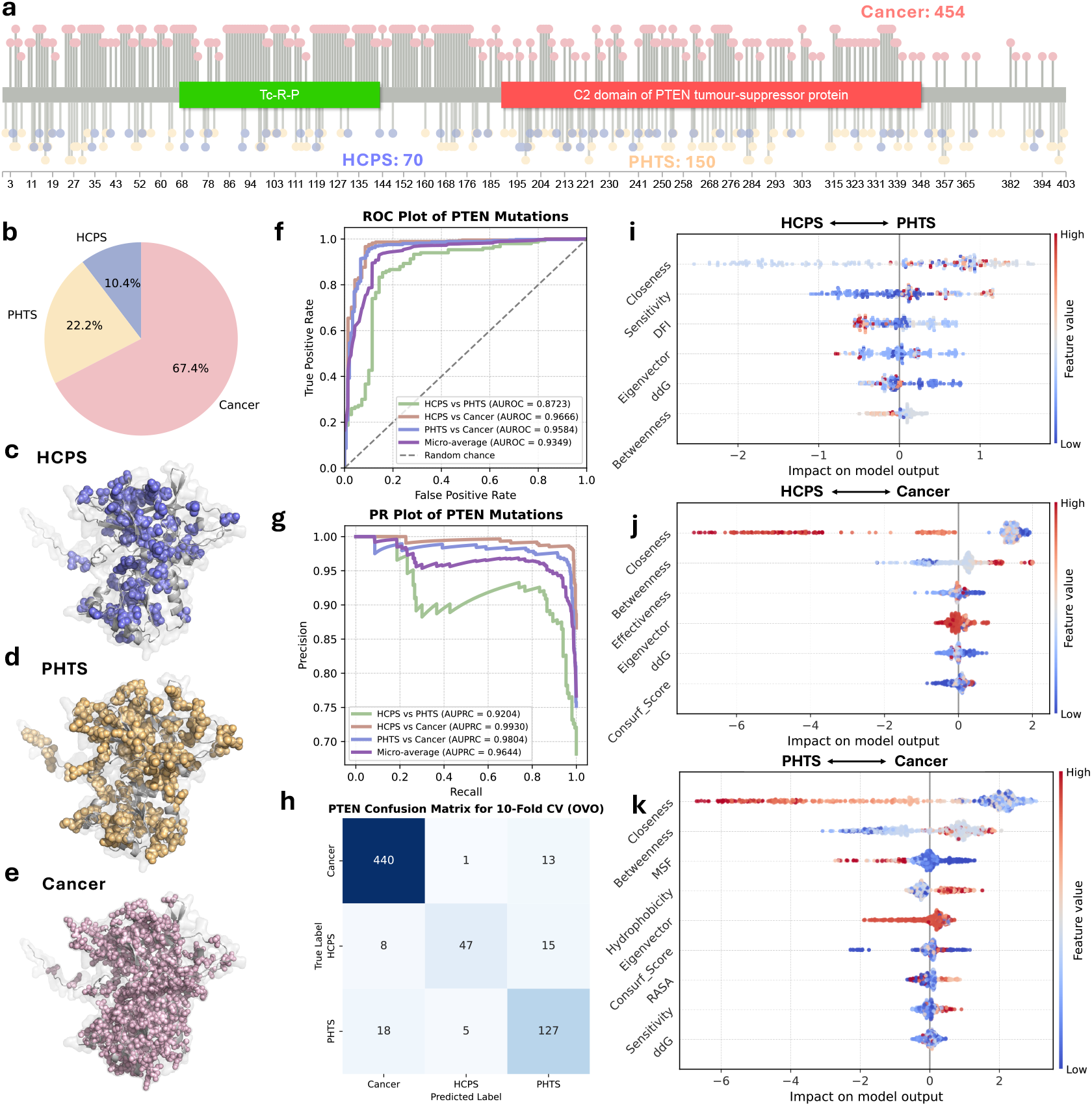
protPheMut classify mutations’ multiple phenotypic effects. a) The lollipop plot shows the distribution of missense mutations on PTEN protein and mutation’s phenotypes. b) The pie chart shows the HCPS, PTHS and cancer variants in dataset. c) The distribution of mutations related to HCPS in PTEN structure. d) The distribution of mutations related to PHTS in PTEN structure. e) The distribution of mutations related to cancer in PTEN structure. f) ROC plot shows the performance of PhenoScore in pairwise comparisons of multiple phenotypes. g) Precision-recall plot of PhenoScore in pairwise comparisons of multiple phenotypes. h) Confusion matrix from 10-fold straight cross-validation using threeclass classifier with One-vs-One strategy on stacking model of XGBoost and LightGBM. i-k) Beeswarm plots show the global and local interpretation of pairwise comparison model using SHAP.

As shown in Figures 3f and 3g, to distinguish PTEN phenotypes, protPheMut performed pairwise comparisons of HCPS, PHTS, and cancer. In the classification tasks for distinguishing between HCPS and PHTS, six key features were used: ΔC, sensitivity, DFI, ΔEC, ΔΔG, and ΔBC. The optimal combination of Gradient Boosting, LightGBM, and CatBoost achieved an AUC of 0.8723 and an AUPRC of 0.9204. For the HCPS vs. cancer classification, also six key features were utilized: ΔC, ΔBC, effectiveness, ΔEC, ΔΔG, and conservation score, resulting in an AUC of 0.9636 and an AUPRC of 0.9930. In the PHTS vs. cancer classification, nine key features were employed: ΔC, ΔBC, MSF, hydrophobicity, ΔEC, conservation score, RASA, sensitivity, and ΔΔG, yielding an AUC of 0.9666 and an AUPRC of 0.9804. The final base models for HCPS vs. cancer and PHTS vs. cancer classification were all Gradient Boosting, XGBoost, and CatBoost. The micro-averaged AUC and AUPRC across all tasks were 0.9349 and 0.9644, respectively.

Performance was higher for distinguishing HCPS or PHTS from cancer but lower for HCPS vs. PHTS, likely due to phenotypic similarities, as both HCPS and PHTS are considered as NDDs. Network topology feature ΔCC consistently emerged as the most critical feature across all classifications. Additional features, such as ΔBC, effectiveness/MSF, and conservation score, were vital for distinguishing HCPS or PHTS from cancer, while sensitivity, ΔEC, and ΔΔG were more significant for HCPS vs. PHTS (Figures 3i-k).

Based on these observations, we constructed a multi-class classification model using a stacking approach with LightGBM and CatBoost as base models and logistic regression as the meta model, employing the One-vs-One (OVO) strategy. The confusion matrix indicated strong performance in classifying cancer and PHTS mutations. However, HCPS mutations were occasionally misclassified as PHTS due to phenotypic overlap (out of 70 HCPS mutations, 47 were correctly classified as HCPS, while 15 were misclassified as PHTS). More detailed analysis of PI3Kα can be found at Supplementary Text S3.

### 3.3 protPheMut web server

Figure 4 illustrates the key components of the protPheMut web platform (http://netprotlab.com/protPheMut), designed to streamline data analysis for users. The platform automates tasks such as feature computation, statistical analysis, machine learning model training, and result interpretation, generating PhenoScores efficiently. Users can visualize results directly on the webpage for quick insights. A detailed user guide is available on the platform.

**Fig. 4.**
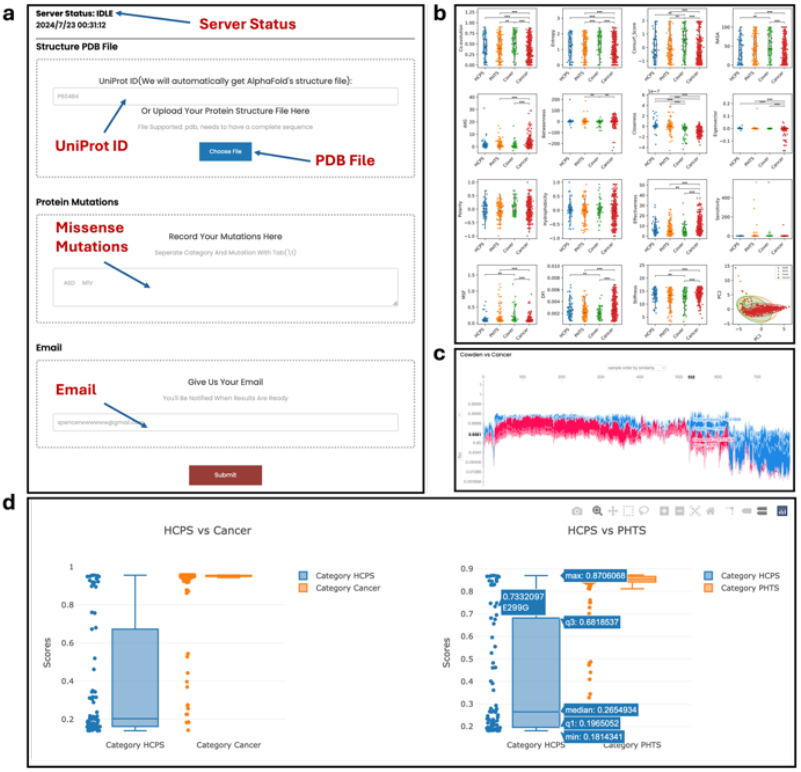
protPheMut web server. a) protPheMut upload form. b) auto feature calculation and statistic tests. c) flow chart of model explanation, Blue represents a negative contribution to the model, red represents a positive contribution to the model, and the graph represents the model’s feature contribution to judging all samples. d) boxplot representing mutations prediction and scoring.

## 4 Discussion

In this study, we developed an interpretable ML framework to assess the phenotypic impacts of human protein missense mutations, with specific applications to PI3Kα and PTEN. Unlike traditional tools that focus on pathogenicity prediction, our work emphasizes phenotype-specific analysis, offering a nuanced perspective on how pathogenic mutations drive distinct disease phenotypes. By introducing the PhenoScore, we provide a novel metric to quantify the phenotypic tendencies of mutations, bridging the gap between prediction and biological interpretation. This approach allows for a deeper understanding of the multi-phenotype effects of pathogenic mutations, particularly in cases of comorbidities between cancer and NDDs, addressing a critical need for unified and biologically informed predictive methods.

protPheMut extends the utility of protein dynamics, which have been shown to improve prediction accuracy in tools like Rhapsody. While Rhapsody (Ponzoni, et al., 2020) leverages ENM, protPheMut incorporates a broader spectrum of features associated with protein missense mutations. These features include alterations in physico-chemical properties, network topology parameters, and protein folding energy. By integrating these additional dimensions of information and leveraging advanced ML techniques, protPheMut provides a more comprehensive and robust framework for analyzing the phenotypic effects of mutations, making it particularly well-suited for complex biological applications. Additionally, protPheMut demonstrates the capability to classify multi-phenotypes, a critical advancement in understanding the complex biological consequences of mutations. By integrating dynamics features, sensitivity or effectiveness, in PTEN mutations, protPheMut captures the nuanced effects of mutations that may drive multiple disease phenotypes. This functionality is particularly valuable for analyzing mutations associated with comorbid conditions, such as NDDs and cancer with more dispersed phenotypes, providing insights into shared and distinct dynamical mechanisms underlying these phenotypes.

Moreover, protPheMut is interpretable. The SHAP-based model explanation provides both global and local interpretations. The global interpretation reveals the importance of all features, while the local interpretation shows the contribution of each feature to the prediction for each specific mutation. Through model explanations, we observe both commonalities and differences in the behavior of various parameters across missense mutations in different proteins. A common finding is that changes in network topology and dynamic parameters play a significant role in differentiating phenotypes. Previous studies have shown that changes in network topology can reflect the allosteric effects of proteins (Kapetis, et al., 2017; Zhao, et al., 2024). Our findings suggest that mutations within the same protein leading to diverse phenotypes are often linked to allosteric mechanisms. Additionally, differences in parameter importance between PI3Kα and PTEN mutations underscore protein-specific mechanisms underlying phenotypic variations. This level of granularity provides actionable insights that could inform targeted therapeutic strategies. For instance, understanding the role of network topology in phenotypic prediction could inspire interventions to modulate these properties, offering opportunities for mutation-specific treatments.

Despite its strengths, the protPheMut framework has certain limitations. The tool’s sensitivity to data volume and unbalanced datasets can affect classification performance, especially when phenotypes are highly correlated. This was evident in the disparity in AUC values observed between PI3Kα and PTEN classification tasks. Additionally, our current study was constrained by the limited scope of analyzed proteins, restricting the generalizability of our findings. Future research aims to expand protPheMut’s application to a broader range of protein mutations. The inherently complex relationship between protein mutations and phenotypes poses challenges for binary classification, particularly for mutations lacking prior information. To address this, we plan to incorporate additional data modalities. For example, integrating epigenomic and genomic features, as demonstrated in recent studies (Liu, et al., 2024), could enhance prediction accuracy. Another promising direction is the inclusion of protein-protein interaction data. Pathway-related interactions could provide a more holistic understanding of mutation effects across multi-protein systems (Xiong, et al., 2024), enabling phenotypic predictions at a network level.

## References

Adzhubei, I., Jordan, D.M. and Sunyaev, S.R. Predicting functional effect of human missense mutations using PolyPhen-2. Curr Protoc Hum Genet 2013;Chapter 7:Unit7 20.

Barozi, V., et al. Revealing SARS-CoV-2 M(pro) mutation cold and hot spots: Dynamic residue network analysis meets machine learning. Comput Struct Biotechnol J 2024;23:3800–3816.

Chen, S., et al. A genomic mutational constraint map using variation in 76,156 human genomes. Nature 2024;625(7993):92–100.

Cheng, J., et al. Accurate proteome-wide missense variant effect prediction with AlphaMissense. Science 2023;381(6664):eadg7492.

Cheng, N., et al. Comparison and integration of computational methods for deleterious synonymous mutation prediction. Brief Bioinform 2020;21(3):970–981.

Elia Venanzi, N.A., et al. Machine Learning Integrating Protein Structure, Sequence, and Dynamics to Predict the Enzyme Activity of Bovine Enterokinase Variants. J Chem Inf Model 2024;64(7):2681–2694.

Gross, A.M., et al. Advancing RAS/RASopathy therapies: An NCI-sponsored intramural and extramural collaboration for the study of RASopathies. Am J Med Genet A 2020;182(4):866–876.

Hopf, T.A., et al. Mutation effects predicted from sequence co-variation. Nat Biotechnol 2017;35(2):128–135.

Jumper, J., et al. Highly accurate protein structure prediction with AlphaFold. Nature 2021;596(7873):583–589.

Kapetis, D., et al. Network topology of NaV1.7 mutations in sodium channel-related painful disorders. BMC Syst Biol 2017;11(1):28.

Landrum, M.J., et al. ClinVar: public archive of relationships among sequence variation and human phenotype. Nucleic Acids Res 2014;42(Database issue):D980-985.

Li, B., et al. De novo mutation of cancer-related genes associates with particular neurodevelopmental disorders. J Mol Med (Berl) 2020;98(12):1701–1712.

Liang, Z., Verkhivker, G.M. and Hu, G. Integration of network models and evolutionary analysis into high-throughput modeling of protein dynamics and allosteric regulation: theory, tools and applications. Brief Bioinform 2020;21(3):815–835.

Liu, Y., et al. MAGPIE: accurate pathogenic prediction for multiple variant types using machine learning approach. Genome Med 2024;16(1):3.

Liu, Y., et al. SHP2 clinical phenotype, cancer, or RASopathies, can be predicted by mutant conformational propensities. Cell Mol Life Sci 2023;81(1):5.

Nussinov, R., et al. Cell phenotypes can be predicted from propensities of protein conformations. Curr Opin Struct Biol 2023;83:102722.

Nussinov, R., Tsai, C.J. and Jang, H. How can same-gene mutations promote both cancer and developmental disorders? Sci Adv 2022;8(2):eabm2059.

Nussinov, R., Tsai, C.J. and Jang, H. Neurodevelopmental disorders, immunity, and cancer are connected. iScience 2022;25(6):104492.

Nussinov, R., et al. Neurodevelopmental disorders, like cancer, are connected to impaired chromatin remodelers, PI3K/mTOR, and PAK1-regulated MAPK. Biophys Rev 2023;15(2):163–181.

Nussinov, R., et al. Review: Cancer and neurodevelopmental disorders: multi-scale reasoning and computational guide. Front Cell Dev Biol 2024;12:1376639.

Orloff, M.S., et al. Germline PIK3CA and AKT1 mutations in Cowden and Cowden-like syndromes. Am J Hum Genet 2013;92(1):76–80.

Pacini, L. and Lesieur, C. A computational methodology to diagnose sequence-variant dynamic perturbations by comparing atomic protein structures. Bioinformatics 2022;38(3):703–709.

Pejaver, V., et al. Inferring the molecular and phenotypic impact of amino acid variants with MutPred2. Nat Commun 2020;11(1):5918.

Pinero, J., et al. The DisGeNET knowledge platform for disease genomics: 2019 update. Nucleic Acids Res 2020;48(D1):D845–D855.

Ponzoni, L. and Bahar, I. Structural dynamics is a determinant of the functional significance of missense variants. Proc Natl Acad Sci U S A 2018;115(16):41644169.

Ponzoni, L., et al. Rhapsody: predicting the pathogenicity of human missense variants. Bioinformatics 2020;36(10):3084–3092.

Portelli, S., et al. Distinguishing between PTEN clinical phenotypes through mutation analysis. Comput Struct Biotechnol J 2021;19:3097–3109.

Shihab, H.A., et al. Predicting the functional, molecular, and phenotypic consequences of amino acid substitutions using hidden Markov models. Hum Mutat 2013;34(1):57–65.

Sim, N.L., et al. SIFT web server: predicting effects of amino acid substitutions on proteins. Nucleic Acids Res 2012;40(Web Server issue):W452–457.

Sondka, Z., et al. COSMIC: a curated database of somatic variants and clinical data for cancer. Nucleic Acids Res 2024;52(D1):D1210–D1217.

Stenson, P.D., et al. The Human Gene Mutation Database (HGMD((R))): optimizing its use in a clinical diagnostic or research setting. Hum Genet 2020;139(10):1197–1207.

Stenson, P.D., et al. The Human Gene Mutation Database: towards a comprehensive repository of inherited mutation data for medical research, genetic diagnosis and next-generation sequencing studies. Hum Genet 2017;136(6):665677.

UniProt, C. UniProt: the Universal Protein Knowledgebase in 2023. Nucleic Acids Res 2023;51(D1):D523–D531.

Varadi, M., et al. AlphaFold Protein Structure Database in 2024: providing structure coverage for over 214 million protein sequences. Nucleic Acids Res 2024;52(D1):D368-D375.

Venot, Q., et al. Targeted therapy in patients with PIK3CA-related overgrowth syndrome. Nature 2018;558(7711):540–546.

Verkhivker, G.M., et al. Allosteric Regulation at the Crossroads of New Technologies: Multiscale Modeling, Networks, and Machine Learning. Front Mol Biosci 2020;7:136.

Xiao, F., et al. Dissecting mutational allosteric effects in alkaline phosphatases associated with different Hypophosphatasia phenotypes: An integrative computational investigation. PLoS Comput Biol 2022;18(3):e1010009.

Xiong, D., et al. A structurally informed human protein-protein interactome reveals proteome-wide perturbations caused by disease mutations. Nat Biotechnol 2024.

Yavuz, B.R., et al. Neurodevelopmental disorders and cancer networks share pathways, but differ in mechanisms, signaling strength, and outcome. NPJ Genom Med 2023;8(1):37.

Yehia, L., Keel, E. and Eng, C. The Clinical Spectrum of PTEN Mutations. Annu Rev Med 2020;71:103–116.

Zhao, Y., et al. Insights into Activation Dynamics and Functional Sites of Inwardly Rectifying Potassium Channel Kir3.2 by an Elastic Network Model Combined with Perturbation Methods. J Phys Chem B 2024;128(6):1360–1370.

Zhu, F., et al. PPICT: an integrated deep neural network for predicting inter-protein PTM cross-talk. Brief Bioinform 2023;24(2).

Zhu, F., et al. Leveraging Protein Dynamics to Identify Functional Phosphorylation Sites using Deep Learning Models. J Chem Inf Model 2022;62(14):3331–3345.

