## Supplementary Text S1-3 for "protPheMut: An Interpretable Machine Learning Tool for Classification of Cancer and Neurodevelopmental Disorders in Human Missense Variants"

### protPheMut Supplementary Text

### S1 protPheMut detailed framework

#### 1.1 Sequenced-based computational analysis

Based on Blastp and multiple sequence alignment (MSA), we calculate the **Shannon information entropy S(i)** and **mutual information MI.** Blastp established local database of protein sequences from the UniProt sequence database UniRef50(Suzek, et al., 2007), evalue is set as 1e-50, and a maximum of 3000 sequences can be searched. The comparison of all sequences takes too long and is not efficient. Meanwhile, it is not necessary to only consider sequences with higher evalue, otherwise it is inaccurate. 200 sequences were randomly selected from a maximum of 3000 sequences with evalue greater than 1e-50. If less than 200 sequences are found, all sequences are compared. The MSA is conducted by Clustal Omega (Sievers, et al., 2011) based on the sequences searched from Blastp (Altschul, et al., 1990). S(i) indicates the stability of amino acid sites in evolution and is calculated using the following formula:

$$S(i) = -\sum_{j=1}^{20} p_j \log(p_j) \tag{1}$$

where p(j) represents the relative frequency of the type j amino acid at position $i^{th}$ in multiple sequence alignment. In multiple sequences, when there is only one amino acid at a position, the Shannon entropy is 0, indicating that the amino acid at the position is determined. When all 20 amino acids appear at this position with superior probability, Shannon entropy reaches the maximum, indicating that the amino acids at this position are completely uncertain. S(i) varies in proteins in the range of 0≤ $S$(i) ≤ 3.0, a lower S(i) indicates a higher conservation. Mutual information (MI) is calculated as a parameter to reflect coevolution between amino acid residues using the following formula:

$$I(i,j) = \sum_{xi=1}^{21} \sum_{yj=1}^{21} P(xi,yj) log \frac{P(xi,yj)}{P(xi)P(yj)} \tag{2}$$

where p(xi, yj) is the joint probability, i and j, of amino acid types x and y observed at sequence positions respectively; p(xi) is the marginal/singlet probability of the x-type amino acid at position i. I(i, j)varies in the range $[0, I_{max}]$in response to completely unrelated and most relevant residue pairs. The co-evolution of mutations was measured by the average MI value corresponding to each residue. The calculation of S(i) and MI is conducted by ProDy (Zhang, et al., 2021). The **conservation** scoring tool Rate4Site (Pupko, et al., 2002) was used to score the MSA results for the conservative type of each amino acid site. The lower the conservative score, the more conservative the site at the sequence level.

#### 1.2 Structure & energy analysis

The Accessible Surface Area (ASA) measures the contact area of a molecule with the solvent and is typically calculated using the Shrake-Rupley algorithm, where a probe ball rolls over the protein surface. The probe ball's radius combines the atomic radius with the solvent radius to determine the contact area. This calculation highlights the differences in ASA between residues buried within the protein and those exposed on the surface. To assess the relative exposure of residues, Relative Accessible Surface Area (RASA) is used. **RASA** normalizes ASA to provide insight into how each residue's exposure compares to its idealized maximum exposure in an extended state. This normalized measure helps in understanding residue accessibility in the context of the protein's folded structure. RASA can be calculated using the following formula:

$$RASA = ASA_{residue}/ASA_{max} \tag{3}$$

where $ASA_{residue}$ is the ASA of the amino acid residue in protein and $ASA_{max}$ is the ASA of the residue in an extended state. The calculation of RASA is based on DSSP (Touw, et al., 2015). The change in the Gibbs energy of folding (**ΔΔG**) is a key index to measure the effect of protein mutation on its stability. The difference in the free energy of the protein before and after the mutation was calculated to assess how the mutation affects the folding and function of the protein. The calculation of the ΔΔG value usually involves a complex energy assessment, including a comprehensive consideration of factors such as interatomic interactions, hydrogen bonding, and hydrophobic effects in the protein structure. We use FoldX5 (Delgado, et al., 2019) for ΔΔG calculation.

#### 1.3 Node-weighted amino acid contact energy network construction

Node-weighted amino acid contact energy network (NACEN) is constructed based on the amino acid contact energy network (Yan, et al., 2020). The function of a protein comes from the high-level structure it folds to form, and the internal residues of the protein interact with each other. In graph theory, the change or collapse of key nodes may lead to global paralysis. In AACEN, proteins are transformed into graph structures, with each amino acid residue as a node and whether there are edges between the nodes depends on the environment-dependent residue contact energy (ERCE) between the residues (Yan, et al., 2018). When the ERCE between two residues is less than 0, the two residues are connected. When the ERCE between two residues is greater than or equal to 0, the two residues are not connected. The ERCE betweenness residue v and u can be calculated using the following formula:

$$ERCE_{vu} = -ln \frac{N_{vu}N_{00}C_{v0}C_{u0}}{N_{v0}N_{u0}C_{vu}C_{00}} \tag{4}$$

where $N_{vu}$ is the number of contacts between amino acid residues i and j observed in the actual protein structure, $N_{00}$ is the number of contacts between two amino acids in a randomized state, $N_{v0}$ and $N_{u0}$ are the total numbers of exposures of amino acids i and j in a particular environment and $C_{v0}$, $C_{u0}$, $C_{vu}$, $C_{00}$ are environmental correction factors that take into account the local environment in which the amino acid is located. The adjacent matrix $\Gamma_{vu}$ of AACEN is:

$$\Gamma_{vu} = \begin{cases} 0, ERCE_{vu} \geq 0 \\ 1, ERCE_{vu} < 0 \end{cases} \tag{5}$$

Based on AACEN, NACEN considers the physical and chemical features of amino acids from the AAindex database (Kawashima, et al., 2008). Since important residues with catalytic function are often distinguishable from other amino acids in terms of hydrophobicity and polarity, and we use these two parameters to weight each amino acid residue. For each mutant protein, we calculated the difference between its parameters and the wild type, including global **Hydrophobicity** changes, **Betweenness** changes, **Closeness** changes, **Eigenvector** changes and **Clustering Coefficient** changes. Betweenness measures how often a node lies on the shortest paths between other pairs of nodes in a network. It reflects the node's importance in network communication; nodes with higher betweenness play a crucial role in information transfer and can act as key bottlenecks in the network.

$$C_B(v) = \sum_{s\neq v\neq t} \frac{\sigma_{st}(v)}{\sigma_{st}} \quad (6)$$

where $\sigma_{st}$ is the number of shortest routes from node s to node t, $\sigma_{st}(v)$ is the number of shortest paths through node v. Closeness measures how close a node is to all other nodes in a network. It reflects the node's efficiency in accessing other nodes and its overall centrality within the network. Nodes with higher closeness can reach other nodes more quickly and are typically more central to the network's structure, thus having greater influence and connectivity.

$$C_C(v) = \frac{n-1}{\sum_{u\neq v} d(v,u)} \quad (7)$$

where $d(v,u)$ is the length of the shortest from node v to node u and n is the number of nodes in network. Eigenvector centrality measures a node's influence by considering not only its connections but also the importance of its neighbors. Nodes with high eigenvector centrality are well-connected to other influential nodes, making them key players in the network.

$$C_E(v) = \frac{1}{\lambda} \sum_{u\in N(v)} A_{vu} x_u \quad (8)$$

where $\lambda$ is the maximum eigenvalue, $N(v)$ is set of neighbor nodes of node v, $A_{vu}$ is the element of node v and u in the adjacency matrix A. The Clustering Coefficient is a measure used in network theory to quantify the degree to which nodes in a graph tend to cluster together. Specifically, it measures how close a node's neighbors are to being a complete clique. For a given node, it is the ratio of the number of actual edges between its neighbors to the maximum possible number of edges that could exist between those neighbors.

$$C(v) = \frac{2T(v)}{deg(v)(deg(v)-1)} \quad (9)$$

Where is the number of edges between the neighbors of node $v$ and $\deg(v)$ is the number of neighbors (degree) of node $v$.

#### 1.4 ENM based dynamic network construction

Protein Elastic Network Models (ENM) are computational tools used to study protein dynamics and function (Ponzoni, et al., 2020). These models include the Gaussian Network Model (GNM) and the Anisotropic Network Model (ANM). GNM simplifies the protein structure into an elastic network of $C_\alpha$ atoms connected by spring $\gamma$ with uniform force constants. This approach effectively predicts low-frequency vibration patterns and provides insights into protein flexibility and stability. The total potential energy of GNM network with n nodes can be calculated using the following formula:

$$V_{GNM} = \frac{\gamma}{2} \sum_{v,u}^{N} \Delta R_v \Gamma_{vu} \Delta R_u \quad (10)$$

where $\gamma$ is the force constant uniform for all springs in the network. $\Delta R_v$ is the three-dimension fluctuation vector as $\Delta X^T = [\Delta X_1, \Delta X_2, \ldots \Delta X_N]$, $\Delta Y^T = [\Delta Y_1, \Delta Y_2, \ldots \Delta Y_N]$ and $\Delta Z^T = [\Delta Z_1, \Delta Z_2, \ldots \Delta Z_N]$. $\Gamma_{vu}$ is the $uv^{th}$ element in the $N \times N$ Kirchoff matrix $\Gamma$ which can be written as:

$$\Gamma_{vu} = \begin{cases} -1, v \neq u \cap R_{vu} \leq r_c^{GNM} \\ 0, v \neq u \cap R_{vu} > r_c^{GNM} \\ -\sum_{v,v\neq u} \Gamma_{vu}, v \neq u \end{cases} \quad (11)$$

where $r_c^{GNM}$ representing cutoff distance in GNM in protPheMut is 7Å. ANM offers a more precise simulation of protein dynamics by accounting for different movement directions. In ANM, the dynamic system can be described by Hessian matrix H written as:

$$H = \begin{pmatrix} H_{11} & \cdots & H_{1N} \\ \vdots & \ddots & \vdots \\ H_{N1} & \cdots & H_{NN} \end{pmatrix} \quad (12)$$

where Hessian matrix contains $N \times N$ elements and each element is a $3 \times 3$ matrix defined as:

$$H_{vu} = \begin{bmatrix} \frac{\partial^2 V_{vu}}{\partial x_v \partial x_u} & \frac{\partial^2 V_{vu}}{\partial x_v \partial y_u} & \frac{\partial^2 V_{vu}}{\partial x_v \partial z_u} \\ \frac{\partial^2 V_{vu}}{\partial y_v \partial x_u} & \frac{\partial^2 V_{vu}}{\partial y_v \partial y_u} & \frac{\partial^2 V_{vu}}{\partial y_v \partial z_u} \\ \frac{\partial^2 V_{vu}}{\partial z_v \partial x_u} & \frac{\partial^2 V_{vu}}{\partial z_v \partial y_u} & \frac{\partial^2 V_{vu}}{\partial z_v \partial z_u} \end{bmatrix} \quad (13)$$

where $V_{vu}$ is harmonic potential calculated by:

$$V_{vu} = \frac{\gamma}{2} (R_{vu} - R_{vu}^0)^2 \quad (14)$$

where $R_{vu}$ and $R_{vu}^0$ are the instantaneous and equilibrium distances between node v and node u in ANM. The dynamic features, **Effectiveness** and **Sensitivity**, are derived from the ANM model by averaging the rows and columns of the Perturbation Response Scanning (PRS) matrix (Atilgan and Atilgan, 2009). This approach involves calculating the average responses of different perturbations across various directions to evaluate how effectively and sensitively a molecular system reacts to structural changes. In dynamic networks, effectiveness refers to a node's efficiency in performing its functions or facilitating information flow, with high-effectiveness nodes often having greater connectivity and playing crucial roles. Sensitivity describes how a node responds to changes or disturbances, with high-sensitivity nodes reacting significantly to minor alterations or disruptions. **Stiffness** is calculated based on ANM by averaging columns of stiffness matrix d defined as:

$$d_{vu} = \sqrt{\frac{k_B T}{\lambda_k} cos\, \alpha_{vu} |\Delta R_v - \Delta R_u|} \quad (15)$$

where $k_B$ is the Boltzmann constant, T is the absolute temperature, $\lambda_k$ is the eigenvalue and cos $\alpha_{vu}$ is the cosine of the angle change after external force is applied. Dynamic Flexibility Index (**DFI**) and Mean Square Fluctuations (**MSF**) are calculated based on GNM where a detailed description of these two features can be found in Bahar's work (Ponzoni, et al., 2020). The computational construction of ENM is based on ProDy (Ponzoni, et al., 2020).

#### 1.5 Statistical analysis and machine learning framework

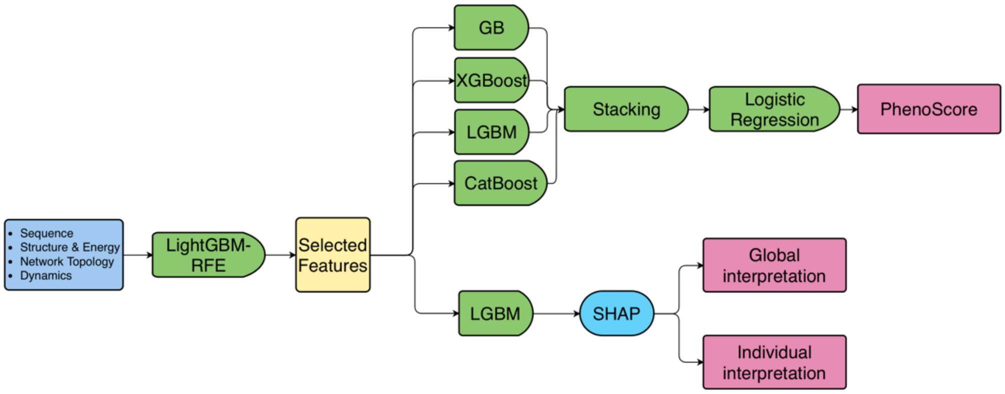

**Fig 1.** Detailed machine learning framework.

Fig 1 shows the diagram of detailed framework of protPheMut machine learning framework and Table 1 shows the mutation features we used in protPheMut. For each phenotype, statistical tests were conducted on fifteen parameters to assess differences. When the sample size was greater than or equal to 30, a t-test was employed. For sample sizes less than 30, the Wilcoxon test was utilized. P-values were calculated to quantify the degree of difference observed. Principal Component Analysis (PCA) was performed on each phenotype's data to reduce dimensionality and reveal key patterns. PCA was executed to extract the principal components explaining the maximum variance, and scatter plots of the first two principal components were generated to visualize the distribution and clustering of phenotypes. For evaluating the discriminative accuracy of each feature, we performed ROC curve analysis. ROC curves were generated by plotting the true positive rate against the false positive rate for each feature. The area under the ROC curve (AUC) was computed to quantify the performance of each feature, with higher AUC values indicating better discriminatory ability. We constructed a correlation matrix for 15 parameters using Spearman's rank correlation coefficient. This non-parametric method measures the strength and direction of the monotonic relationship between two variables. The Spearman correlation coefficient is calculated based on the ranked values of the data rather than the raw values:

$$\rho = 1 - \frac{6\sum_{i=1}^{n} d_i^2}{n(n^2-1)} \quad (16)$$

where $d_i^2$ is the difference between the ranks of each pair of observations and $n$ is the number of paired observations.

**Table 1.** The mutation features we use in protPheMut.

| Sequence | Biophysical | Network | Dynamics |
|---|---|---|---|
| Entropy | Hydrophobicity | Betweenness | Effectiveness |
| Coevolution | rASA | Closeness | Sensitivity |
| Consurf_Score | ΔΔG | Eigenvector | Stiffness |
| | | Clustering Coefficient | DFI |
| | | | MSF |

Multiple perspective features show different mutations' information. Sequence information reveals mutation conservation, while energy and structural changes indicate variations in protein folding and free energy. Amino acid interactions network analysis reflect how mutations affect global protein structure. Residues in dynamic network with low stability impact local structures, whereas highly stable sites transmit changes to more distant regions, showcasing allosteric effects. Machine learning will be used to integrate these data into a comprehensive model of protein mutations. We began by using Recursive Feature Elimination (RFE) with LightGBM (n_estimators=1000, max_depth=5, learning_rate = 0.01, weighted = "balanced") to evaluate 15 parameters. This process involved recursively training the model, ranking feature importance, and eliminating less significant features until only one feature remained. We assessed the selected features using the mean accuracy of stratified 5-fold cross-validation to identify the best parameter combination and contain information from all aspects.

In our analysis, we employed Tree SHapley Additive exPlanations (SHAP) to facilitate both global and individual interpretability of our model (Lundberg and Lee, 2017). SHAP values are grounded in cooperative game theory, where they derive from Shapley values which is a method originally developed to allocate contributions among players in a cooperative game. In the context of machine learning, SHAP provides a robust framework for interpreting model predictions by attributing the contribution of each feature to the final prediction. This method allows us to quantify the global importance of parameters, highlighting their overall impact on the model, while also offering detailed insights into the contribution of each feature for individual samples. By leveraging SHAP, we gain a comprehensive understanding of how different features influence predictions at both aggregate and granular levels.

Additionally, we employed a Stacking Model, an ensemble learning technique, to enhance performance and generalization. This method integrates predictions from multiple base models through a meta-model, which in our case was a logistic regression model. Logistic regression, suitable for binary classification, combines the output probabilities from the base models to produce a final prediction which can be defined as:

$$\begin{cases} \hat{y} = \sigma(z) = \frac{1}{1+e^{-z}} \\ z = \omega_1 x_1 + \omega_2 x_2 + \cdots + \omega_n x_n + b \end{cases} \quad (17)$$

where z represents the output of the linear model, $\sigma(z)$ is a logical function used to map the output of the linear model to the interval (0, 1), representing the probability of the positive class of the sample. We tested combinations of four machine learning algorithms: GradientBoost, XGBoost, LightGBM, and CatBoost. With grid search to get best parameters, we used stratified 10-fold straight cross-validation prediction to generate probabilities. The meta-model aggregated these probabilities, and we evaluated the model performance using area under ROC curve (AUC) to determine the best model combination and its score. We used this method to obtain the integrated probability of each mutation in the dataset for the binary comparison of phenotypes, defining this as the PhenoScore, which represents the score of the mutation in terms of the phenotype, used to measure the phenotypic effects of the mutation.

We use these statistical metrics to measure PhenoScore and other scores generated by other tools: AUC (Area Under ROC Curve), AUPRC (Area Under Precision-Recall Curve), Accuracy, Precision, Recall, F1-Score, and MCC (Matthews Correlation Coefficient). These metrics provide a comprehensive evaluation of model performance, considering various aspects of binary classification, including distinguishing power (AUC and AUPRC), overall correctness (Accuracy), and the balance between positive predictive power and sensitivity (Precision, Recall, F1-Score). Additionally, MCC offers a balanced measure accounting for true and false predictions. The statistical metrics are calculated using the following formulas:

$$\text{Accuracy} = \frac{\text{TP} + \text{TN}}{\text{TP} + \text{TN} + \text{FP} + \text{FN}} \quad (18)$$

$$\text{Precision} = \frac{\text{TP}}{\text{TP} + \text{FP}} \quad (19)$$

$$\text{Recall} = \frac{\text{TP}}{\text{TP} + \text{FN}} \quad (20)$$

$$\text{F1-Score} = \frac{2 \times \text{Precision} \times \text{Recall}}{\text{Precision} + \text{Recall}} \quad (21)$$

$$\text{MCC} = \frac{(\text{TP} \times \text{TN}) - (\text{FP} \times \text{FN})}{\sqrt{(\text{TP} + \text{FP})(\text{TP} + \text{FN})(\text{TN} + \text{FP})(\text{TN} + \text{FN})}} \quad (22)$$

where TP is True Positives, TN is True Negatives, FP is False Positives, and FN is False Negatives.

In addressing multi-phenotype multi-class classification problems, we employ the One-vs-One (OVO) strategy. OVO strategy involves breaking down a multi-class problem into multiple binary classification problems, where each pair of classes is compared. For the core machine learning strategy, we build a Stacking Model by combining models that have shown strong performance in binary classifications. The feature parameters used are a combination of features that exhibit high importance across the binary classification systems. To evaluate the model, we apply 10-fold straight cross-validation, generating a confusion matrix to assess performance. The machine learning framework is based on Python package scikit-learn (version 1.3.0), the data analysis is based on Python (version 3.10) and R (version 4.3.2) language and plots are generated using the Python package matplotlib (version 3.7.3) and seaborn (version 0.12.2).

### S2 Oncoprotein PI3Kα detailed analysis

#### 2.1 Features Generation

The features related to PI3Kα mutations are generated automatically by protPheMut. Table 2 shows the Pvalue conducted by T-test and AUC in single feature ROC analysis in comparing cowden syndrome group and cancer group, the features are significant features in cancer and Cowden syndrome comparison. Fig 2 shows the classification ability of features without machine learning models (single feature ROC curve) and 15 features correlation. Through feature correlation, we observed that features in sequence conservation (Co-evolution, Entropy and Conservation score) and features in inter-molecular dynamics (Effectiveness, Sensitivity, MSF, DFI and Stiffness) are respectively highly correlated, which shows these features may reveal similar information in PI3Kα protein missense mutations. It's also worth noting that rASA is highly correlated with dynamic features.

**Table 2.** Features with significant P Value (P Value < 0.05) in cancer and Cowden syndrome comparison. Number in red color shows the most significant feature in T-test or in ROC analysis.

| Feature | P Value | AUC |
|---|---|---|
| Consurf_Score | **1.061e-06** | 0.6285 |
| ΔΔG | 6.610e-05 | 0.6052 |
| ΔC | 0.0002982 | **0.6824** |
| ΔBC | 0.01127 | 0.5254 |
| Hydrophobicity | 0.01432 | 0.5931 |
| Sensitivity | 0.04200 | 0.5526 |

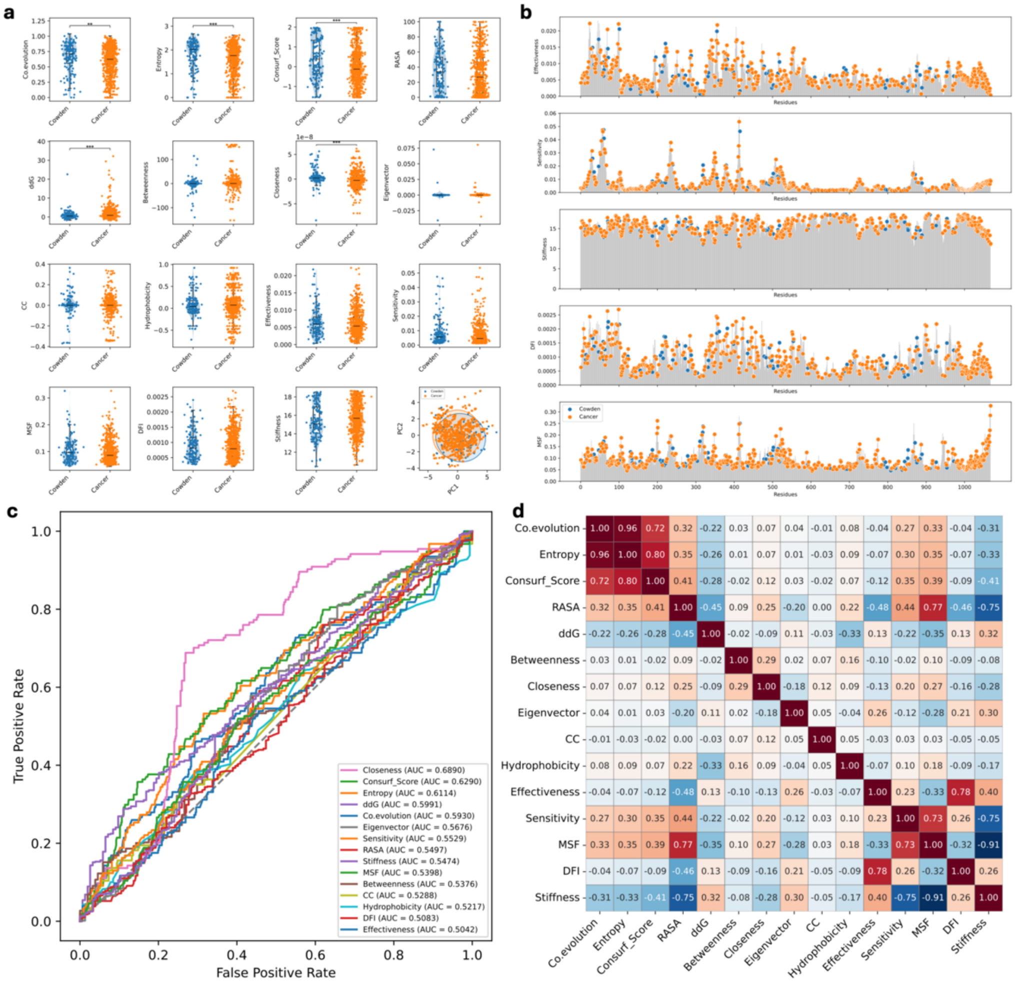

**Fig 2.** Features generated by protPheMut. a) Features visualizations and tests. b) Distribution of dynamic features in sequence. c) Single feature ROC curves with their AUC. d) Features correlation heatmap using Spearman correlation.

#### 2.2 Model Construction Detail

Fig 3a illustrates the process of feature selection using Recursive Feature Elimination (RFE), where the optimal subset of features is identified based on five-fold cross-validation scores. Fig 3b shows the search results within the stacking model, presenting the AUC results corresponding to all model combinations. Fig 4 presents additional model interpretation results using a force plot.

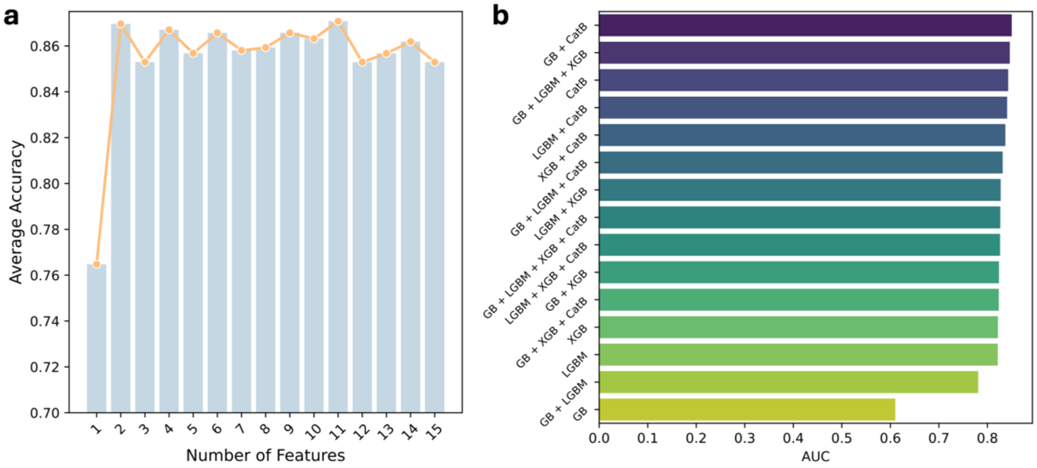

**Fig 3.** Feature selection and stacking model optimization. a) The feature number generated by RFE and 5-fold cross-validation score. b) The AUC of all model combinations as base model in stacking model.

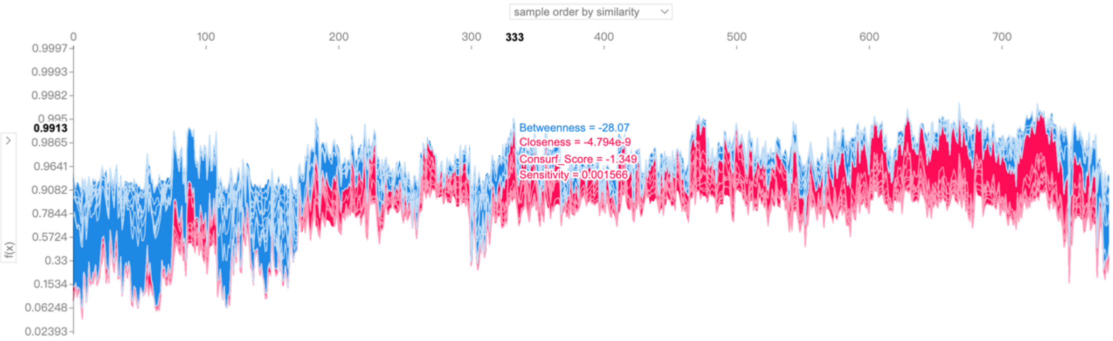

**Fig 4. The force plot of model explanation.** The figure shows SHAP results, visualizing sample contributions to model predictions ordered by similarity. The x-axis represents sample indices, and the y-axis shows SHAP values. Blue and red curves depict feature contributions, which blue is negative and red is positive. The annotation highlights one sample with its decision features, providing insights into model behavior and key sample characteristics.

#### 2.3 Other Evaluation Metrics

In the main analysis, we evaluated the performance of PhenoScore and other benchmark tools using AUC and AUPRC. Table 3 shows the other statistical evaluation metrics of PhenoScore and other methods. These metrics assess the discriminative power and the balance between precision and recall, particularly relevant for imbalanced datasets. PhenoScore achieved the highest AUC and AUPRC values, indicating its superior ability to distinguish between classes and maintain a high precision-recall trade-off compared to other methods.

PhenoScore demonstrated excellent performance with the highest Recall (0.963), F1-Score (0.912), and MCC (0.456), reflecting its strong sensitivity and balanced prediction capability. Its Accuracy (0.850) also surpassed all other tools, underlining its robustness. Benchmark tools such as AlphaMissense and Rhapsody showed competitive Precision values (0.896 and 0.888, respectively) but had lower Recall and F1-Score, highlighting a trade-off between precision and sensitivity. Other methods like EVMutation and FATHMM achieved lower scores across most metrics, indicating limited performance. These findings support PhenoScore's effectiveness in accurately and reliably predicting outcomes, surpassing the performance of current benchmark tools.

**Table 3.** Other statistical metrics to measure performance.

| Method | Accuracy | Precision | Recall | F1-Score | MCC |
|---|---|---|---|---|---|
| PhenoScore | **0.850** | 0.866 | **0.963** | **0.912** | **0.456** |
| AlphaMissense | 0.645 | **0.896** | 0.632 | 0.741 | 0.267 |
| EVMutation | 0.796 | 0.667 | 0.051 | 0.094 | 0.143 |
| PolyPhen-2 | 0.661 | 0.867 | 0.683 | 0.764 | 0.21 |
| Rhapsody | 0.629 | 0.888 | 0.616 | 0.727 | 0.237 |
| MutPred2 | 0.776 | 0.834 | 0.901 | 0.866 | 0.197 |
| FATHMM | 0.735 | 0.257 | 0.182 | 0.213 | 0.061 |
| SIFT | 0.681 | 0.244 | 0.288 | 0.264 | 0.062 |

### S3 Oncoprotein PTEN detailed analysis

#### 3.1 Features Generation

Table 4 shows the Pvalue conducted by T-test and AUC in single feature ROC analysis, the features are significant features in cancer and Cowden syndrome comparison. Fig 5 and Fig 6 shows visualization and some basic analysis of the features.

**Table 4.** Features with significant P Value (P Value < 0.05) in HCPS, PTHS, and cancer comparisons. Number in red color shows the most significant feature in T-test or in ROC analysis.

| Group | Feature | P Value | AUC |
|---|---|---|---|
| **HCPS vs. PHTS** | | | |
| | MSF | **0.01467** | 0.5587 |
| | Stiffness | 0.02298 | **0.5740** |
| **HCPS vs. Cancer** | | | |
| | Coevolution | 2.780e-07 | 0.6562 |
| | Entropy | 8.462e-07 | 0.6524 |
| | Conservation score | 1.728e-05 | 0.6129 |
| | rASA | 1.288e-07 | 0.6019 |
| | ΔΔG | 0.0003510 | 0.5775 |
| | ΔC | **2.814e-25** | **0.9196** |
| | ΔEC | 4.234e-05 | 0.6244 |
| | Effectiveness | 3.251e-08 | 0.5914 |
| | MSF | 1.313e-05 | 0.6186 |
| | DFI | 2.751e-10 | 0.6012 |
| | Stiffness | 2.629e-08 | 0.5962 |
| **PHTS vs. Cancer** | | | |
| | Coevolution | 1.859e-05 | 0.6394 |
| | Entropy | 3.020e-05 | 0.6324 |
| | Conservation score | 0.006954 | 0.6264 |
| | rASA | 0.01523 | 0.6474 |
| | ΔC | **3.110e-17** | **0.8783** |
| | ΔEC | 3.616e-06 | 0.5919 |
| | Effectiveness | 0.006430 | 0.6557 |
| | DFI | 0.01185 | 0.6401 |
| | Stiffness | 0.01460 | 0.6563 |

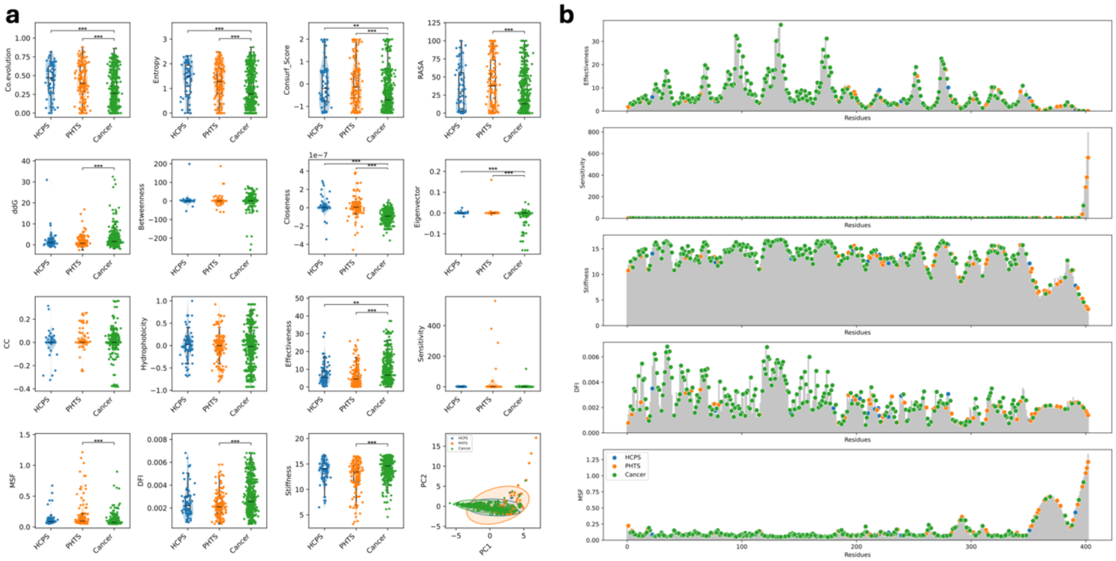

**Fig 5.** Features generated by protPheMut. a) Features visualizations and tests. b) Distribution of dynamic features in sequence.

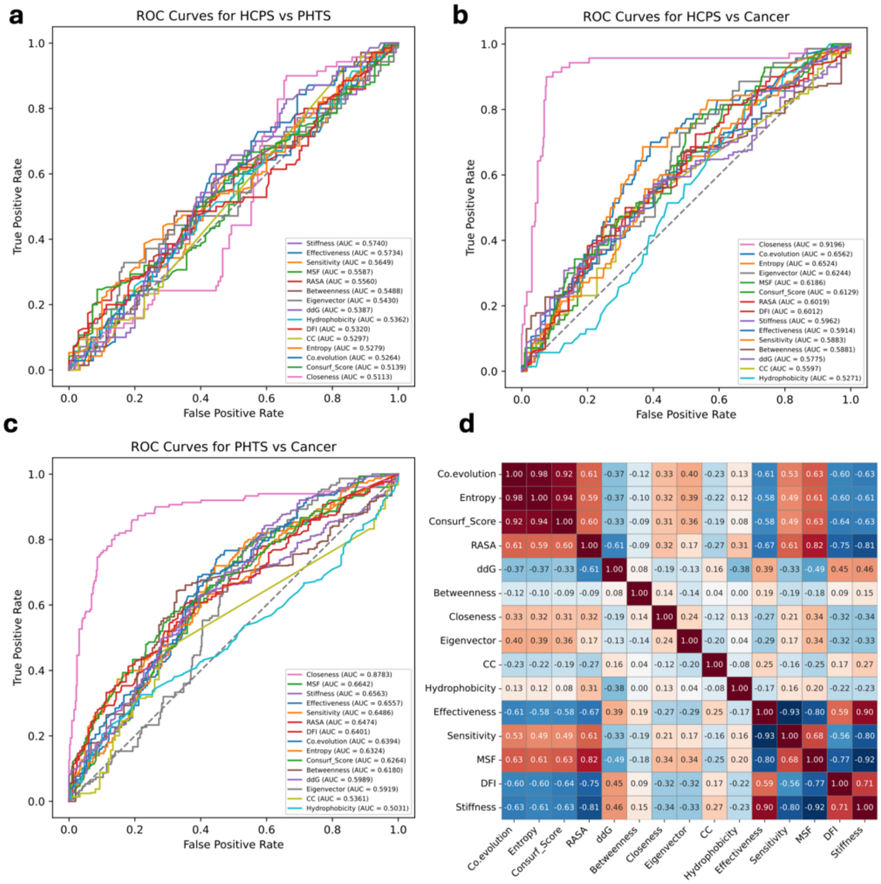

**Fig 6.** Features generated by protPheMut. a-c) Single feature ROC curves with their AUC. d) Spearman correlation heatmap.

#### 3.2 Model Construction Detail

Fig 7 shows the model construction detail and SHAP force plots of PTEN mutations in HCPS, PHTS, and cancer comparisions.

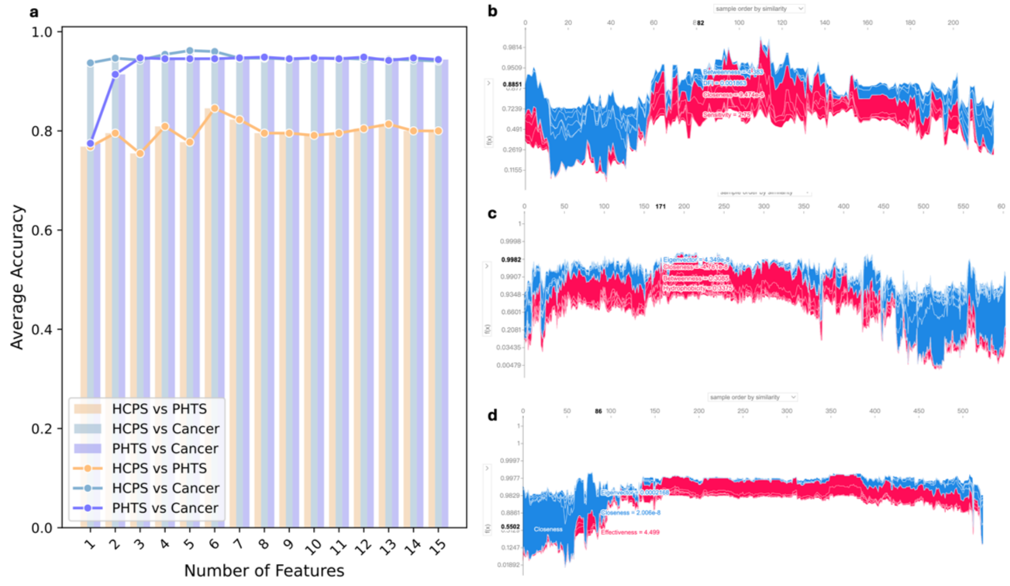

**Fig 7.** PTEN model construction detail and explanation. a) The feature number generated by RFE and 5-fold cross-validation score in HCPS, PHTS, and cancer comparisons. b) Force plot of the model explanation in HCPS vs. PHTS. c) Force plot of the model explanation in HCPS vs. cancer. d) Force plot of the model explanation in PTHS vs. cancer.

##### 3.3 Other Evaluations

**Table 5.** Other statistical metrics to measure performance.

| Group | Accuracy | Precision | Recall | F1-Score | MCC |
|---|---|---|---|---|---|
| HCPS vs. PHTS | 0.855 | 0.878 | 0.913 | 0.895 | 0.658 |
| HCPS vs. Cancer | 0.940 | 0.948 | 0.974 | 0.961 | 0.837 |
| PHTS vs. Cancer | 0.958 | 0.966 | 0.987 | 0.976 | 0.810 |

#### Supplementary Reference

Altschul, S.F., *et al.* Basic local alignment search tool. *J Mol Biol* 1990;215(3):403-410.

Atilgan, C. and Atilgan, A.R. Perturbation-response scanning reveals ligand entry-exit mechanisms of ferric binding protein. *PLoS Comput Biol* 2009;5(10):e1000544.

Delgado, J., *et al.* FoldX 5.0: working with RNA, small molecules and a new graphical interface. *Bioinformatics* 2019;35(20):4168-4169.

Kawashima, S., *et al.* AAindex: amino acid index database, progress report 2008. *Nucleic Acids Res* 2008;36(Database issue):D202-205.

Lundberg, S. and Lee, S.-I. A Unified Approach to Interpreting Model Predictions. In.; 2017. p. arXiv:1705.07874.

Ponzoni, L., *et al.* Rhapsody: predicting the pathogenicity of human missense variants. *Bioinformatics* 2020;36(10):3084-3092.

Pupko, T., *et al.* Rate4Site: an algorithmic tool for the identification of functional regions in proteins by surface mapping of evolutionary determinants within their homologues. *Bioinformatics* 2002;18 Suppl 1:S71-77.

Sievers, F., *et al.* Fast, scalable generation of high-quality protein multiple sequence alignments using Clustal Omega. *Mol Syst Biol* 2011;7:539.

Suzek, B.E., *et al.* UniRef: comprehensive and non-redundant UniProt reference clusters. *Bioinformatics* 2007;23(10):1282-1288.

Touw, W.G., *et al.* A series of PDB-related databanks for everyday needs. *Nucleic Acids Res* 2015;43(Database issue):D364-368.

Yan, W., *et al.* Node-Weighted Amino Acid Network Strategy for Characterization and Identification of Protein Functional Residues. *J Chem Inf Model* 2018;58(9):2024-2032.

Yan, W., *et al.* ANCA: A Web Server for Amino Acid Networks Construction and Analysis. *Front Mol Biosci* 2020;7:582702.

Zhang, S., *et al.* ProDy 2.0: increased scale and scope after 10 years of protein dynamics modelling with Python. *Bioinformatics* 2021;37(20):3657-3659.
